# Following cell type transitions in space and time by combining live-cell tracking and endpoint cell identity in intestinal organoids

**DOI:** 10.1101/2022.06.27.497728

**Authors:** Xuan Zheng, Max A. Betjes, Yvonne J. Goos, Guizela Huelsz-Prince, Hans Clevers, Jeroen S. van Zon, Sander J. Tans

**Affiliations:** AMOLF, Amsterdam, Netherlands; Hubrecht Institute, Royal Netherlands Academy of Arts and Sciences (KNAW) and UMC Utrecht, 3584 CT Utrecht, the Netherlands; Bionanoscience Department, Kavli Institute of Nanoscience Delft, Delft University of Technology, Delft, Netherlands

## Abstract

Elucidating the dynamics of cellular differentiation in space and time is key to advancing organoid biology and technology. Direct visualization of differentiation patterns is challenging, however, in part because of the difficulty of simultaneously detecting all relevant cell types by fluorescence imaging. Here we present TypeTracker, which determines the differentiation trajectories of all cells within a region of interest, including their type transitions, growth, divisions, and changes in spatial organization. We show how the lineage tree topology, as determined by automated cell tracking, allows for the retrospective identification of cell types across multiple generations. The data revealed various surprising aspects of intestinal organoid differentiation, including symmetric fate adoption by sister cells, cell type abundance regulated by division, and type commitment of cells occurring prior to their spatial reorganization. Our method can be applied broadly to study other organoid systems and to screen for compounds that affect differentiation programs.

## Introduction

The past decade has brought groundbreaking developments in organoid technology^1-3^. Key to the potential of organoids is their ability to *in vitro* generate the diverse cell types and 3D architectures that characterize *in vivo* organ tissue. Hence, organoids enable a host of new approaches to study organ development, self-renewal, and pathology^4^. Various features of the cellular architecture of organoids have been established using immunostaining, fluorescence microscopy, and single-cell RNA sequencing^5-7^. However, these methods do not detect the dynamics of cellular differentiation or spatial organization, and hence are limited in their ability to identify underlying patterns, phenotypes, and mechanisms. Lineage tracing^8-10^ can visualize the offspring of certain cells in a snapshot image but does not identify cell type transitions or characterize cell movement, thus leaving the spatio-temporal differentiation dynamics largely unaddressed.

In parallel, important advances have been achieved in 3D video microscopy. Techniques such as confocal and light-sheet imaging have visualized the dynamics of cell proliferation and collective cell migration in major developmental transitions^11-13^. Machine learning has more recently enabled automated tracking of individual cells in time, in developing embryos as well as in diverse organoid systems^13-16^. Yet, video microscopy has far been limited in studying spatiotemporal differentiation programs. Cell fate markers can in principle be followed using fluorescent protein markers^17-19^. However, the significant number of cell types (over 5 in intestinal organoids) and the need to label the nuclei for cell tracking, presents major challenges in terms of phototoxicity, genetic construction, and wavelength overlap. Indeed, while fluorescence microscopy can visualize the Lgr5 intestinal stem cell marker^20^, it has been challenging to expand this approach to include the other intestinal cell types^17^, and to combine it with cell tracking to reveal the spatio-temporal differentiation dynamics, which indeed remains uncharacterized. This capability is crucial, as key open questions about the formation and maintenance of multi-cellular architectures involve the moments and locations that cells commit to a new fate, the genealogical relations between cell types, and the neighboring cells that can define dynamically changing spatial niches^21^. These issues are ideally addressed by characterizing most if not all cells in a particular region of interest. Such a method would be of direct relevance to many types of organoid systems, as well as drug screening applications.

Here we present TypeTracker, an approach that integrates cell type identification with cell tracking to directly view the organoid differentiation dynamics. It works by propagating the differentiation state of each cell back in time along the branched lineage tree, using cell tracking to determine the lineage tree during organoid growth, and several rounds of antibody staining to determine the endpoint cell types (**Fig. 1a**). The resulting multi-dimensional data identifies the moments that cells commit to a new type, their movements and genealogical relations to all other cells, and what neighbors they have interacted with at any point in time. It can be used generally for any organoid system with proliferating and differentiating cells, and by most laboratories, as it requires only confocal microscopy, immunostaining, and software for data integration.

**Fig. 1.**
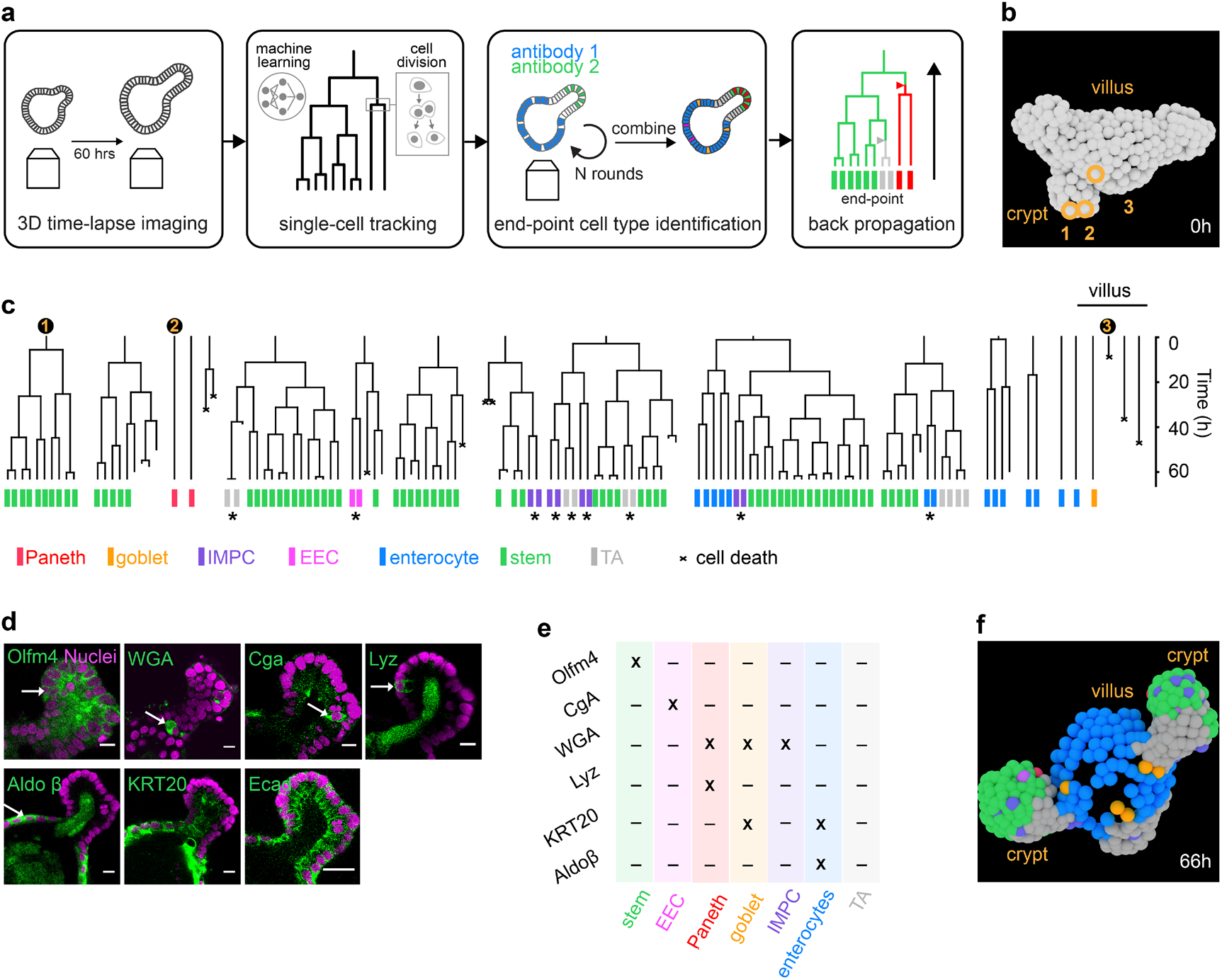
TypeTracker method overview and primary data. **a**, Method workflow. Step 1: Image several organoids in 3D for 60h. Step 2: Track all cells and determine lineage tree using convolutional neural network. Step 3: Identify cell type at movie endpoint using multiple rounds of antibody staining. Step 4: Map cell types onto lineage tree, and infer historical cell types including type transitions (triangles). **b**, Reconstructed cell positions at start of lineage tree. **c**, Example lineage trees and their mapped endpoint cell types. Numbers: lineages of three cells seen in panel b. Stars: endpoint sister pairs of same type, but of different type than cousins, suggesting type transition in mother cell. **d**, Nuclear marker (magenta) and antibody or dye staining (green): Olfactomedin 4 (Olfm4), Wheat Germ Agglutinin (WGA), Chromogranin A(Cga), Lysozyme (Lyz), Aldolase β (Aldoβ), Cytokeratin 20 (Krt20) and Ecadherin (Ecad). **e**, Cell type identification table. **f**, Reconstructed cell positions and type at end of the lineage tree (panel c), 66 hours after the start (panel b).

We applied TypeTracker to mouse intestinal organoids, which are 3D cellular assemblies that display crypt and villus domains (**Fig. 1b)**. Our data showed that differentiation was controlled to a surprising degree by information transmitted through the lineage. Specifically, sister cells systematically adopted the same fate, including secretory fates, with their fate being determined by the mother cell. The lineage trees showed significant variability, with some sub-lineages moving up the crypt-villus axis while other related lineages in the same tree remained deep within the crypt in the stem cell zone. The order of events was notable. We did not observe the lineages first moving up and away from their relatives, and then committing to a new type, as expected in current models. Rather, we found lineages that committed to a new type first, while still nearby their relatives that retained stem cell fate and then separated from them subsequently. These data are consistent with cells sorting in space following specification. Our TypeTracker approach can be used broadly to elucidate differentiation programs of organoid systems, how they are modulated by disease mutations or external cues, and can be readily combined with fluorescence-based monitoring of processes such as WNT signaling, apoptosis, or actin polymerization.

## Results

### Overview of TypeTracker

Our method consists of four main steps (**Fig. 1a**). First, movies of organoids growing in Basement Membrane Extract (BME) gel are recorded by 3D confocal microscopy (**Extended Data Fig. 1a-c**). We focus on intestinal organoids and image complete organoids or their crypt protrusions every 12 minutes for 60 hours, using a H2B-mCherry marker to detect cell nuclei (**Extended Data Fig. 1d**). Second, we employ a convolutional neural network^14^ to track the nuclei in space and time across several generations (**Fig. 1b, Extended Data Fig. 1e**), and visualize their genealogical relations using lineage trees (**Fig. 1c, Extended Data Fig. 2**). Consistently, proliferative lineages, which display up to seven divisions, are mostly found at the crypt bottom, while non-dividing cells are abundant in the villus-like region of the organoid. Third, we perform multiple rounds of antibody staining to determine the cell types at the movie endpoint, as detailed below **(Fig. 1d, e)**. Hence, we identify the main types found in the intestine: stem, enteroendocrine, Paneth, goblet, enterocyte, transit-amplifying, as well as an immature mucus producing type. Fourth, we infer the cell types at earlier timepoints, starting at the endpoint of the lineage tree and progressively moving back along each branch to the beginning, using a set of rules that we describe in detail below. Hence, one obtains virtual organoids that present the spatial location, division events, movements, and type of all cells (**Fig. 1f**). The completeness of this method, which identifies all cells in a region of interest and associated lineage trees, enables our backpropagation approach and analysis of spatio-temporal correlations.

### Cell type identification

We developed an approach to map cell type data, as obtained by several rounds of antibody staining, onto the endpoint of the movie and lineage trees (**Fig. 1d, e**). This method includes washing protocols that minimize organoid deformations, antibody stripping between rounds, and a min-cost flow solver algorithm^22^ combined with manual correction to link cells in the staining images to the cells at the movie endpoint. Stem cells were identified by Olfactomedin 4 (Olfm4)^23^, enterocytes by Aldolase β (AldoB)^24^ and enteroendocrine cells (EECs) by Chromogranin A (Cga)^25^. Consistently, Olfm4+ cells were found only at the crypt bottom, Cga+ cells in both the crypt and villus regions and AldoB+ cells exclusively in the villus region (**Fig. 1f**). Stem cells and EECs did not stain for any of the other markers used, while enterocytes also stained for Cytokeratin 20 (KRT20)^26^, as expected. Wheat Germ Agglutinin (WGA), which stains intestinal mucus^27,28^, labeled cells in the crypt and villus regions. A subset of these were also positive for Lysozyme (Lyz) and thus identified as Paneth cells^29^, while another subset expressed KRT20 and identified as goblet cells^26^. Consistently, the former often contained granules that are typical of Paneth cells, while the latter had a cup-shaped morphology that characterizes goblet cells (**Extended Data Fig. 3)**. A remaining subset of WGA+ cells expressed neither Lyz nor KRT20. Evidence provided below suggests that these are immature Paneth and goblet cells, which we thus refer to as immature mucus producing cells (IMPCs). We identified a population of cells within the crypt that did not stain for any of our markers, which are identified as transit-amplifying (TA) cells based on evidence discussed below.

### Sister cells adopt the same type

Prominent in our data is that endpoint types generally came in pairs, with sister cells consistently displaying the same fate (**Fig. 1c**). Statistical analysis showed that 97% of all endpoint sister (N=869 sister pairs in 9 organoids) were of the same type.

All types, including secretory types, exhibited this symmetry, as evidenced by the dominant diagonal in the sister-sister cell type histogram (**Fig. 2a**). This consistent type-symmetry between sisters is notable. Intestinal stem cells are proposed to exit from their cell cycle when committing to a secretory fate^30,31^, which would rather yield asymmetry, as one sister may then adopt a secretory fate that the other does not. External cues can, in principle, produce a local environment that is similar for the two sister cells, and hence drive them to the same fate. However, a priori there is then no reason why only sisters should be impacted, as also non-sister cells like cousins can be neighbors. Importantly, the fates of many sister pairs were different than their direct cousins (**Fig. 1c**, stars). This showed that the underlying type transitions did affect sisters specifically and occurred during the observed growth period of the lineage. These data indicated that the type transitions rather occurred in the mother (or earlier generations), which subsequently divided to produce two daughters of the same type. Larger subtrees, which for instance showed two or more sister pairs of the same type (**Fig. 1c**, blue enterocyte subtree, **Extended Data Fig. 2**), were consistent with divisions occurring after type commitment and hence producing cells of the same type.

**Fig. 2.**
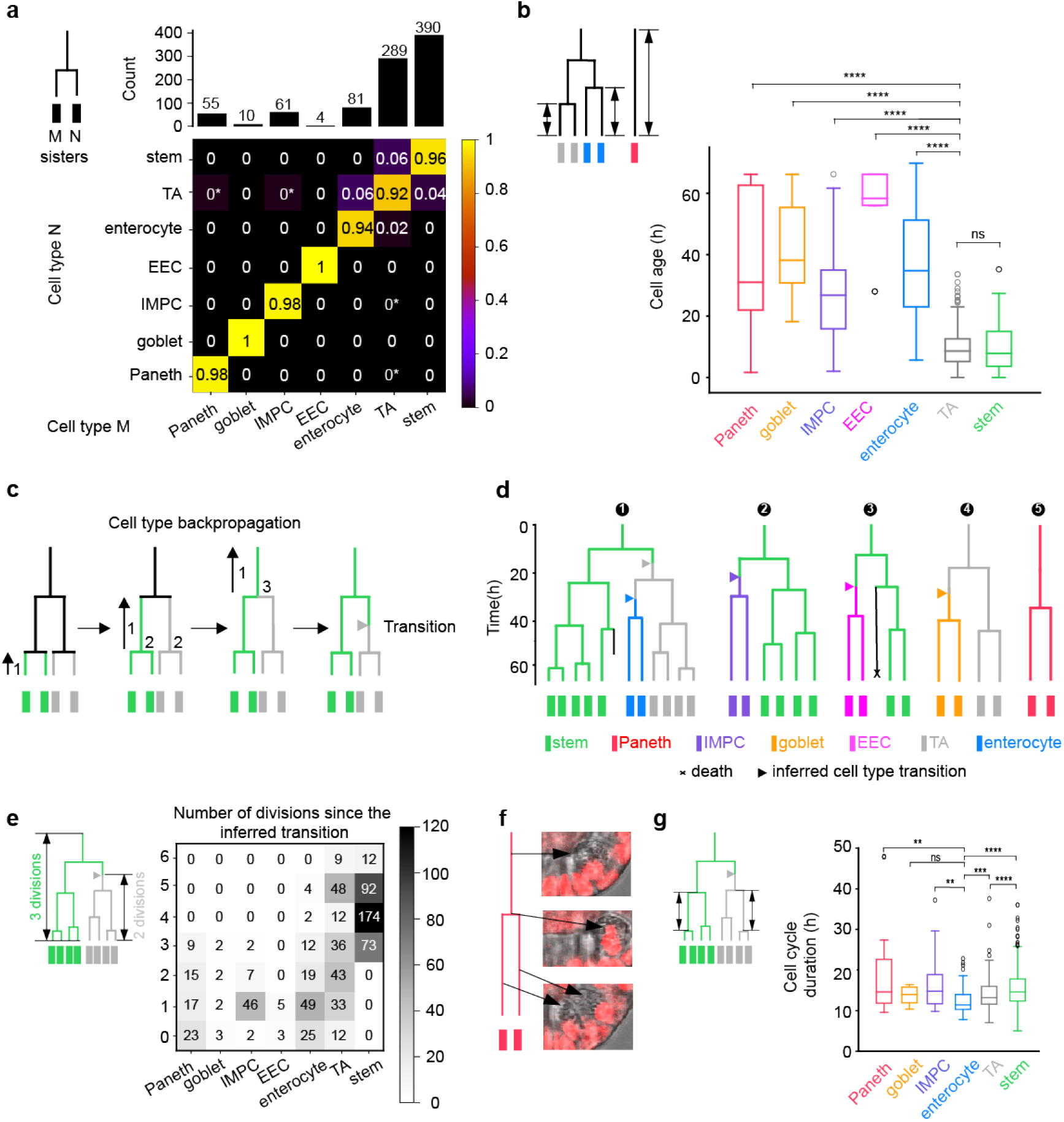
Differentiation pathways and cell type commitment. **a**, 2D histogram indicating the two cell types of sisters at the movie endpoint, as determined using antibody staining. Shown is the number of times a cell type combination was observed divided by the total of each column, indicated on top. Star: low frequency resulting from a single observation. The dominant diagonal shows sisters typically have the same type. **b**, Age of stained cells, quantified as the time since the last division, or since movie start for lineages without division. ****, p ≤ 0.0001; ***, p ≤ 0.001; **, p ≤ 0.01; *, p ≤ 0.05; ns, p > 0.05. TA cells show the same age as stem cells, indicating their proliferative nature. Consistently, the differentiated cells (EECs, enterocytes, Paneth and goblet cells) are older. **c**, Cell-type backpropagation rules. 1: working backwards along each lineage, the inferred type is unchanged when not encountering branchpoints (divisions). 2: for branchpoints, if both daughters are the same type, the mother is assigned that type. 3: If only one daughter a stem cell, the mother is assigned as a stem cell. 4 (not shown): If only one daughter is a TA cell, and the other is not a stem cell, the mother is assigned as a TA cell. In cases 3 and 4, a type-transition is inferred in one daughter (triangle). **d**, Example lineage trees as determined by TypeTracker. **e**, Number of observed divisions for each cell type. **f**, Paneth cell divisions. Granules are seen before and after division, evidencing Paneth cell division. **g**, Cell cycle duration for different cell types. Cell cycle durations were notably similar between types, while enterocytes exhibited lower mean cell cycle durations than TA cells, stem cells and the secretory cell types. All data in this figure were from nine organoids.

### Dependence of cell age on differentiation state

The age of the cells at the movie endpoint was defined as the time since the last observed division (**Fig. 2b**). Notably, the TA cells showed a statistically identical age distribution as the stem cells, in line with the notion that they are highly proliferative^31^. EECs, enterocytes, Paneth, and goblet cells also showed a broad distribution, but with an average age that was substantially larger than the stem and TA cells, consistent with their terminally differentiated nature. The ages of these terminally differentiated cells were rarely below 20 hours (**Fig. 2b**), suggesting that the involved markers (Cga, AldoB, Lyz and KRT20) took a substantial amount of time to become observable.

### Cell type backpropagation

To infer the cell type along the lineages of the family tree, we used the lineage tree topology and the above observations (**Fig. 2a, b**). Starting at the lineage endpoints, we propagated the endpoint types back in time using this process: 1) From one timepoint to the previous timepoint, the type is initially assigned as unchanged if branch points (divisions) are not traversed, but may later be updated when a cell type transition is identified. 2) At branch points, the two involved daughters are considered: if both display the same (inferred) type, the mother is assigned as that type, as discussed above (**Fig. 2a**). 3) If one daughter is a stem cell and the other is not, the mother is assigned as a stem cell, based on the notion that stem cells are generated from stem cells. 4) If one daughter is a TA cell and the other is not, nor a stem cell, the mother is assigned as a TA cell. This rule is based on the TA cells being proliferative like stem cells (**Fig. 2b**), which makes the alternative (the mother has the identity of the non-TA daughter) unlikely. In the last two cases (3 and 4), a type-transition is inferred in one of the daughters. We mark the latter with an arrow halfway the cell cycle **(Fig. 2c)**. These rules allowed the backpropagation of complete trees from the end to the beginning (**Fig. 2d, Extended Data Fig. 4**).

Next, we estimated stemness by quantifying the fluorescence intensity of Lgr5-GFP, a well-known stem-cell marker^32^, in order to further test our method. Note that phototoxicity, which is stronger for GFP than for the mCherry that is used for nuclei tracking, limits the frequency and duration of such cell fate quantification during growth. The GFP signal at the movie endpoint was indeed correlated quantitatively with the measured Olfm4 intensity (**Extended Data Fig. 5a**,**b**). Cells inferred as stem cells by backpropagation indeed also showed higher GFP expression, while inferred enterocytes or TA and goblet cells consistently showed lower GFP (**Extended Data Fig. 5c**). Lineages that were inferred to lose stemness and transition to the TA type indeed showed decreasing GFP (**Extended Data Fig. 5d**). Paneth cell granules, which can be visualized during organoid growth, also showed consistency between inferred and real-time measured type (**Fig. 2f**). Note that limitations of our method are described in the discussion.

### Differentiation pathways

The observed differentiation pathways indicated notable features (**Fig. 2d, Extended Data Fig. 4**). Trees typically displayed 1 to 3 cell type transitions, thus yielding sub-trees of the new type, while the old type was maintained in another sub-tree, and some lineages showed two consecutive type transitions. For instance, a stem cell tree was first shown to spawn a TA-subtree, which in turn generated an enterocyte subtree (**Fig. 2d**, tree 1). This order, in which TA is an intermediate type between stem and terminally differentiated types^33^, is consistent with the sister-sister cell type histogram: besides the dominant diagonal, low-frequency off-diagonal entries indicated stem-TA sister pairs, and a few pairs of one TA and one terminally differentiated type, but never sister pairs showing two different terminally differentiated types (**Fig. 2a**). The latter would be in line with models where stem cells first differentiate into secretory precursors, which in turn can generate different terminally differentiated types. To further probe this notion, we extracted the largest possible subtrees from our data that contained two types, and found that none had two different secretory types, nor one secretory and one absorptive type. Instead, they typically combined TA and either stem (52.2%) or a terminally differentiated state (31.4%) (**Extended Data Fig. 6**), consistent with the sister relations (**Fig. 2a**). Among these subtrees, we also found cases (16.4%) that combined stem and terminally differentiated states (**Fig. 2d**, tree 2 and 3). The latter could indicate that the intermediate TA state lasted for less than one cell cycle and could not be detected, or that these lineages have a negligible TA role.

### Division rather than differentiation rates control enterocyte abundance

Cell type transitions to enterocytes were less frequent than to the secretory types combined (about 1.5-fold), even as enterocytes outnumbered secretory cells. To investigate this issue, we quantified the number of consecutive divisions in each differentiation state (**Fig. 2e**). The EEC, goblet and IMPC secretory states mostly showed one division, sometimes two. In contrast with common thinking^31^, we found Paneth cells to divide as well (**Fig. 2e**). The inferred Paneth cell divisions were indeed confirmed by the continuous presence of granules that are specific to Paneth cells and are observable without labeling (**Fig. 2f**). Next, we quantified the cell cycle duration for each cell type as the time between birth and division. Notably, these complete cell cycles that also showed division at the end were similarly long for all types including secretory and stem types (**Fig. 2g**). After the divisions, the secretory lineages were observed to stop dividing, consistent with a cell cycle exit.

Enterocytes displayed significantly more divisions than secretory cells. The former reached to up to 5 divisions, not far off from the 6 divisions seen for stem and TA states (**Fig. 2e**). Hence, after specification, absorptive lineages generated substantially more cells than secretory lineages, which thus offset their comparatively low transition rate. This finding is consistent with recent data on 2D intestinal enteroid monolayer, in which absorptive lineages showed larger clone sizes than secretory lineages^34^. Enterocytes were even found to exhibit the lowest mean cell cycle duration (12.5 h), significantly lower than for both stem (15.5 h) and TA cells (14.1 h) (**Fig. 2g**). Overall, these results indicate that relative abundance is controlled by the cell lineage dynamics, in particular the number of divisions after specification.

### The spatio-temporal differentiation program

Access to spatial dynamics is a key benefit of our approach. The positions of inferred cell types mapped along the crypt-villus axis as expected, with stem and Paneth most towards the bottom, followed by IMPC, EEC, TA, enterocyte, and goblet^33^ (**Fig. 3a, b, Extended Data Fig. 7a**). In addition, we can now map the point of commitment onto the spatial organoid structure. Commitments to secretory types were broadly distributed along the crypt-villus axis, but typically occurred deeper in the crypt than absorptive commitments (**Fig. 3c**). The latter also distributed broadly and were positioned well within the crypt. Notably, when we mapped full lineage trees along the crypt-villus axis, we found that these committing cells are still in the same approximate spatial region as their stem-cell relatives that had not differentiated (**Fig. 3d, e, Extended Data Fig. 7b, c**). The commitment to secretory and absorptive fates thus occurred early, rather than while moving upwards in the crypt as often assumed^33^. Spatial separation from these stem-cell relatives did occur, after the type transition (**Fig. 3d, e**). This separation was most evident for lineages destined for the villus region (enterocytes and goblet cells), with the stem lineages they arose from remaining confined to the crypt (**Fig. 3d, e**, top trees). Such separation from relatives was not always observed, specifically for cells committing to a secretory type (IMPC) near the crypt bottom and remaining there (**Fig. 3d** bottom tree, **Fig. 3e**, top tree). Consistently, the mean migration speed is highest for enterocytes and goblet cells, and lowest for secretory fates like Paneth cells and IMPC’s (**Fig. 3f**).

**Fig. 3.**
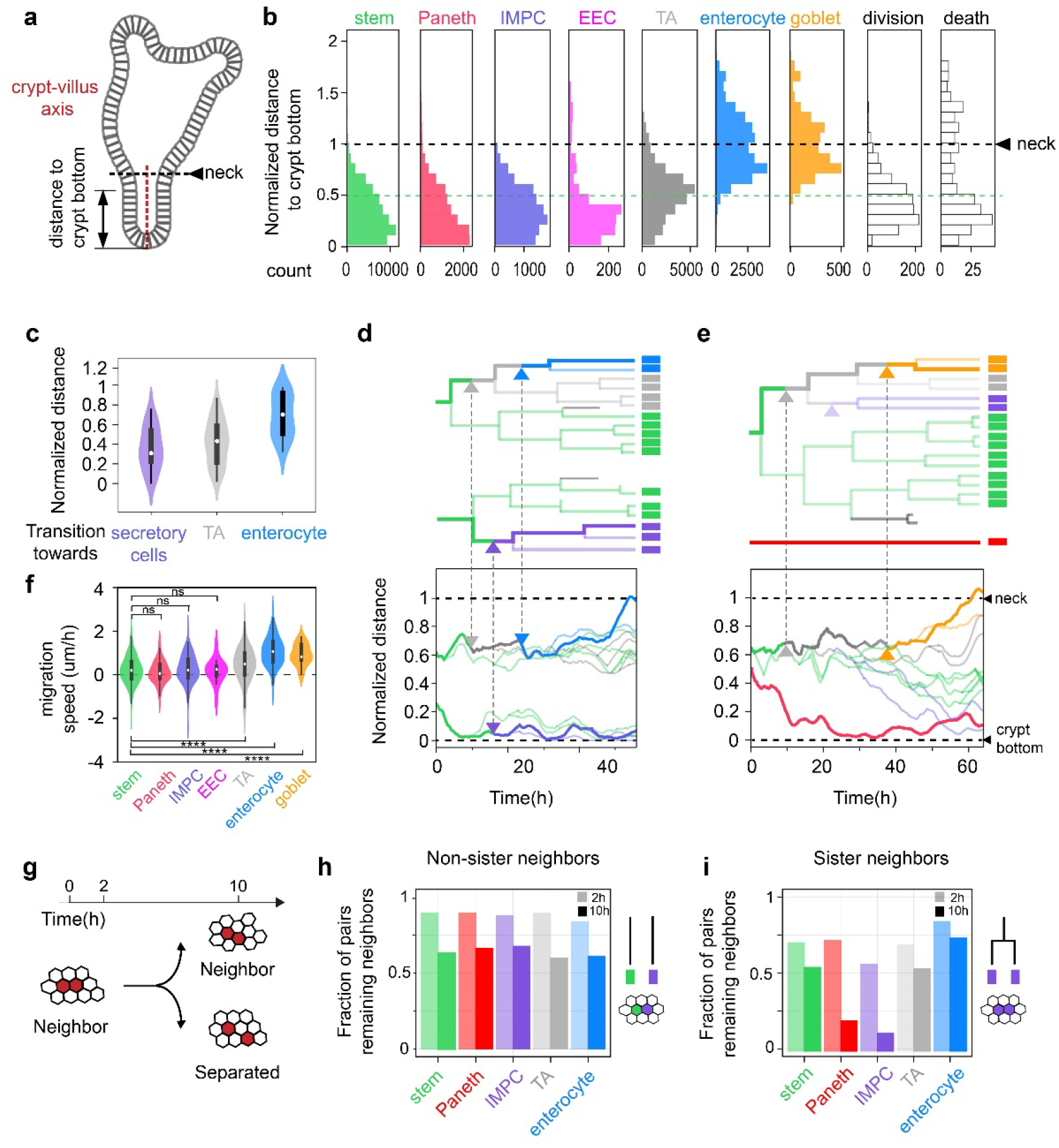
Spatio-temporal differentiation program. **a**, Organoid schematic diagram. **b**, Inferred cell types mapped along the normalized crypt-villus axis. **c**, Position of cells when they commit to a new type. **d and e**, Lineage trees mapped in space along the crypt-villus axis. Bottom trees: Transition to IMPC and position of Paneth cell deep in the crypt surrounded by stem cells. Top trees: Transitions to enterocytes or goblet cells higher in the crypt, but close to their stem cell relatives, followed by their movement to the villus region, rather than the other way around. **f**, Cell migration speed, as quantified by the migration distance along the crypt-villus axis after commitment or movie start, divided by the migration time. **g**, Diagram showing that cell separation is quantified as the fraction of neighboring cells that remain so after 2 and 10 h. **h**, Cell separation of neighbors. This neighbor loss can be quantified for each cell type. Most neighbors remained neighbors even after 10h, independently of cell type. **i**, Cell separation of sisters. Here the sisters generally are of the same type, due to the fate symmetry of sisters. Sisters thus separate faster than non-sisters, in particular sister Paneth cells and IMPCs.

### Promotion of sister separation

Secretory types were typically born as sisters (**Fig. 2a**) and, hence, as neighbors. We wondered how this finding relates to the observation of single Paneth and goblet cells being surrounded by stem cells and enterocytes, respectively^1,35^. In our data, most neighbors that were *not* sisters (and hence can be of different type), were still neighbors after 2h (85%) and 10h (57%), independently of each neighbor’s cell type (**Fig. 3g,h, Extended Data Fig. 7d**). Neighbors that *were* sisters, which almost always were of the same type, separated more frequently after 2h (71% remains neighbors), likely due to arrangements directly following divisions^36^. Surprisingly, Paneth and IMPC sisters showed even stronger separation over longer timescales, and more than the other types, with less than 20% still neighbors after 10h (**Fig. 3i**). Sisters, and in particular of these secretory types, thus appear to rearrange more strongly to achieve interspersion. These findings extend ‘conveyor belt’ models where movement is driven by pushing from growing and dividing neighbors, and instead indicates that the promotion of cell separation contributes to cell type patterning in the epithelium.

## Discussion

While our approach allows visualization of the spatio-temporal differentiation dynamics, it also has a number of limitations. First, cell type transitions may be missed when they are too frequent. Specifically, it cannot deduce the presence of stem or TA cells, or transitions from them, if the tree endpoint only shows differentiated types. This issue can be mitigated partly by limiting the growth duration. Second, if cells would exhibit multiple sequential transitions without dividing, or show reversible transitions back to the stem or TA type, the method would be less suitable. Third, used antibodies may not identify immature types. Long type maturation is indeed suggested by our data, including the dominant sister symmetry (**Fig. 2a**), and the high age of differentiated cells at movie endpoint (**Fig. 2b**), noting that rapid type maturation after commitment would yield sister asymmetry in the staining. This issue impacts marker-based methods more generally. Moreover, our method provides a key advantage in overcoming this limitation, by detecting early commitment using lineage relations even when the phenotype is not detectably expressed yet.

The spatio-temporal view provided by TypeTracker revealed a number of notable findings, even as many observations agree with existing models, like the presence of stem cell and TA zones, and overall migration along the crypt-villus axis (**Fig. 3a, d-f**). Fate commitment is often described to occur in the TA zone, as the cells move up the crypt and down the spatial WNT gradient, with secretory and absorptive progenitors emerging in close vicinity, such that these two fates and their ratios can be regulated by lateral Notch inhibition between neighbors, and with newly specified Paneth cells subsequently moving down to the crypt bottom^33,37^.

Our data showed committing cells positioned at similar locations as their stem cell relatives, and spatial separation taking place subsequently (**Fig. 3c-e**). The latter was clear in particular for absorptive and secretory types that moved towards the villus region, while their stem cell relatives within the same lineage tree remained at the same height. We found commitment even deeper in the crypt for secretory types such as Paneth cells that ended up at the crypt bottom, and hence did not show the proposed downward migration after commitment^33,37^. We did not observe the opposite: stem cells that first move away from their stem cell relatives and then differentiate while continuing to move upwards to the villus region, or downward to the crypt bottom. Overall, our data suggested a picture in which cells first differentiate and subsequently spatially sort and reorganize.

Notable is also the highly symmetric adoption of identical fates by sister cells, including secretory fates. The data indicated that this symmetry stems from genealogical rather than spatial factors, with mother cells committing and passing the new fate down to their daughters (**Fig. 2a-d, 3d, e**). Neighbor analysis showed sister cells to spatially separate even faster than non-sister neighbors (**Fig. 3g-i**). In addition, the majority of the spatially isolated secretory cell pairs (two nearby cells of the same type that are surrounded by cells of different types) were sisters (82%). Importantly, these findings do not suggest or imply that spatial cues are not critical to the mother’s commitment. Yet, our findings contrasts with secretory fate commitment in the intestine being associated with cell cycle exit^38^, and raises the question whether at least one division is important to completing differentiation, as suggested^39^. Interestingly, previously reported clonal expansion of enterocytes and Goblet cells after fate commitment in mouse colon are consistent with our findings^10^. Our data further showed that the ratio between absorptive and secretory cells was affected more by increased divisions after enterocyte specification than by more frequent enterocyte specification events (**Fig. 2e**), as for instance controlled by Notch signaling^40^. Notch signaling between neighbors is proposed to restrict secretory fate to a single cell^40^. Our observation of neighboring sisters with the same secretory fate implies that Notch signaling occurs early and transiently, likely in the mothers of secretory cells.

Our method can be applied broadly to study the spatio-temporal differentiation programs of organoid systems, and how they are impacted by external conditions such as metabolic compounds or interleukins, interacting cell types including bacteria and immune cells, and disease mutations. Its focus on spatial and temporal characterization makes it distinct from and complementary to other methods such as single cell RNA sequencing or multiplexed tissue imaging^41,42^. The TypeTracker approach follows systematically all cells in a region of interest, thus allowing direct correlative analysis, and is straightforward to implement as it requires only confocal microscopy, antibody staining, and the algorithms we present here. It can also readily be combined with other measurements, such as endpoint Single Molecule Fluorescence In Situ Hybridization (smFISH) and real-time fluorescence microscopy of various cellular processes and signals.

## Author contributions

X.Z., H.C., J.V.Z. and S.T. conceived the research. X.Z., J.V.Z. and S.T. wrote the manuscript with the input and discussion of all authors. X.Z., Y.J.G, G.H.P tested the antibodies and dyes and optimized protocols of fixation. Y.J.G adapted the antibody stripping protocol. X.Z. carried out the time-lapse imaging of organoids, followed by multiple rounds of antibody staining. X.Z. and Y.J.G conducted live-cell tracking, endpoint cell type identification and mapping cell types to lineages. X.Z. developed the Python scripts to perform most of the data analysis. M.A.B. developed the Python scripts to analyze cell neighbors and separation rates.

## Acknowledgements

We thank N. Sachs and J. Beumer for providing the mouse intestinal organoids, R. Kok, K. Spoelstra, P. Ender for detailed discussions about the project and comments of the manuscript, S. Semrau and K. Ganzinger for comments and critical reading of the manuscript. X.Z. was funded by the NWO Building Blocks of Life grant from the Dutch Research Council, number 737.016.009. M.A.B. was funded by the NWO Groot grant from the Dutch Research Council, number 2019.085.

