## Supplementary material for "Following cell type transitions in space and time by combining live-cell tracking and endpoint cell identity in intestinal organoids": Methods and Supplementary figures

**Organoid culture.** Murine intestinal organoids carrying both a H2B-mCherry reporter and a Lgr5-GFP reporter were gifts from Norman Sachs and Joep Beumer, from the group of Hans Clevers in Hubrecht Institute. Organoids were embedded in 'domes' of basement membrane extract (BME, Trevingen) in tissue culture plates. They were further submerged in growth medium consisting of murine recombinant epidermal growth factor (EGF 50 ng/ml, Life Technologies), murine recombinant Noggin (100 ng/ml, Peprotech), human recombinant R-spondin 1 (500 ng/ml, Peprotech), n-Acetylcysteine (1 mM, Sigma-Aldrich), N2 supplement (1x, Life Technologies) and B27 supplement (1x, Life Technologies), Glutamax (2 mM, Life Technologies), HEPES (10 mM, Life Technologies), Penicilin/Streptomycin (100 U/ml 100 µg/ml, Life Technologies) in Advanced DMEM/F-12 (Life Technologies). Organoids were kept in incubators at 37 °C and with 5 % $CO_2$. The medium was changed every two days. Each week, organoids were passaged by mechanically dissociating crypts using a narrowed glass pipette.

**Organoid sample preparation for imaging.** In conventional culture conditions, organoids were embedded in 'domes' of BME droplets and were thus at different heights relative to the plate bottom (**Extended Data Fig. 1a**). Imaging organoids located far from the plate bottom required long working distance objectives and increased light exposure, leading to excessive phototoxicity. To improve the imaging procedures, we used the 4 well chambered cover glass (# 1.5 high performance cover glass) from Cellvis as imaging plates. Organoids were broken into single crypts, seeded in the imaging plates and put in fridge (~ 4 °C) for ~ 10 minutes, allowing them to sink downwards to the cover glass. Afterwards, they would be incubated at 37 °C with 5 % $CO_2$ for 20 minutes so that the gel could solidify with organoids settled at the bottom of the wells (**Extended Data Fig. 1b**). Growth medium was added after the incubation. Organoids were then kept in the incubator for around 2 days until the imaging experiments.

**Time-lapse imaging with 3D confocal microscope**. Time-lapse imaging was performed with a scanning confocal microscope (Leica TCS SP8) with a 40 x water immersion objective (NA = 1.10). Experiments were performed at 37 °C and 5 % $CO_2$. More than 20 organoids with already budded crypts were selected for imaging. Stacks of ~ 30 z-slices with 2 µm step size were taken every 12 minutes per organoid. At each timepoint, imaging of H2B-mCherry was conducted with an excitation laser of 552 nm at 1 % of the laser power and the emission signals were collected with Leica HyD hybrid detectors whose filter range was set to be 557 nm - 789 nm.

**Live-cell tracking**. Live-cell tracking was conducted by OrganoidTracker, a software developed by our group[1]. The positions of each nucleus were predicted with a trained neural network and cells were then automatically linked between frames based on the relative positions and nuclear sizes. The software could report warnings when the linking was less reliable and allowed for manual corrections.

**3D reconstruction**. 3D reconstruction of organoids were made with Blender, a free and open-source 3D computer graphics software. Each cell was represented by a 3D sphere and could be colored based on the (inferred) cell types (**Fig. 1b,f**).

**Organoid fixation and permeabilization**. Organoid samples were fixed with 4 % formaldehyde (Sigma-Aldrich) at room temperature. In order to get rid of the gel but keep the organoids attached to the plate, we optimized the fixation protocol. After adding formaldehyde, we waited for ~ 10 minutes and then gentling washed the sample with PBS to remove the gel, which otherwise would hinder the penetration of antibodies and reduce imaging quality. 10 minutes was the optimized waiting time to ensure gel removal and more than 50 % of the imaged organoids attached to the cover glass (**Extended Data Fig. 1c**). After gel removal, organoid samples were incubated in formaldehyde again for 20 minutes to complete the fixation procedures. Following fixation, permeabilization was performed by incubating the samples in 0.2 % Triton-X-100 (Sigma-Aldrich) for one hour at room temperature. All the washing procedures were performed gently to avoid removing organoids from the cover glass.

**Staining with antibodies and dyes**. Following fixation and permeabilization, organoids were blocked with 5 % skim milk in TBS at room temperature for one hour. Subsequently, organoids were incubated in blocking buffer containing primary antibody (see section antibodies) for two days at 4 °C, and then incubated with secondary antibody (see section antibodies) at room temperature for one hour. These procedures would be repeated for each antibody. Regarding the dyes, organoids were incubated with Wheat germ agglutinin (WGA) conjugated to CF©488A (5 ug/ml Biotium) at room temperature for two hours and with RedDot™1 Far-Red Nuclear stain (1 : 200 Biotium) or SYTOX™ Orange Nucleic Acid Stain (1 : 5000 Thermo Fisher Scientific #S 11368) at room temperature for 20 minutes.

**Antibody stripping**. After imaging the results from each round of antibody staining, the primary antibodies were removed by incubation with elution buffer at room temperature for 15 minutes while shaking[2]. This was repeated six times with the elution buffer replaced between consecutive cycles. The elution buffer was prepared by adding 0.5 M Glycine (Sigma-Aldrich), 5 M Urea (Sigma-Aldrich), 5 M Guanidinium chloride (Sigma-Aldrich), 70 mM TCEP-HCL (Sigma-Aldrich) to $H_2O$, with pH adjusted to 2.5.

**Antibodies and dyes**

| **Primary antibodies** | **Dilution** | **Product information** |
|---|---|---|
| Rabbit anti-lysozyme [EC 3.2.1.17] | 1 : 800 | Dako #A0099 |
| Rabbit anti-Olfm4 [D6Y5A] XP® | 1 : 500 | Cell signaling technology #39141 |
| Recombinant rabbit anti-Aldolase B + Aldolase C [EPR3138Y] | 1 : 300 | Abcam #ab75751 |
| Mouse anti-Human Cytokeratin 20 [Clone Ks20.8] | 1 : 500 | Dako #M701929-2 |
| Mouse anti-Chr-A [C-12] | 1 : 50 | Santa Cruz Biotechnology #sc-393941 |
| Rat anti-E-cadherin [DECMA-1] | 1 : 400 | Santa Cruz Biotechnology #sc-59778 |

| **Dyes** | **Dilution** | **Product information** |
|---|---|---|
| WGA conjugated to CF©488A | 5 ug/ml | Biotium |
| RedDot™1 Far-Red Nuclear stain | 1 : 200 | Biotium |
| SYTOX™ Orange Nucleic Acid Stain | 1 : 5000 | Thermo Fisher Scientific #S11368 |

| **Secondary antibodies** | **Dilution** | **Product information** |
|---|---|---|
| Goat anti-rabbit IgG H&L (Alexa Fluor©405) pre-adsorbed | 1 : 1000 | Abcam #ab175654 |
| Goat anti-Rat IgG H&L (Alexa Fluor©555) pre-adsorbed | 1 : 1000 | Abcam #ab150166 |
| Donkey anti-Mouse IgG H&L (Alexa Fluor©647) | 1 : 500 | Thermo Fisher #A31571 |
| Donkey anti-Rabbit IgG H&L (Alexa Fluor©405) pre-adsorbed | 1 : 1000 | Abcam #ab175649 |

The order of staining, optimized to ensure good staining quality for all cell types, was based on the staining quality and stripping difficulty of each antibody (**Extended Data Fig. 1f**).

**Cell type identification**. Olfactomedin 4 (Olfm4) and Chromogranin A (Cga) stained stem and enteroendocrine cells (EECs) respectively[3,4]. Paneth and goblet cells were both stained by Wheat Germ Agglutinin (WGA), which stains mucus[5], and could be distinguished by affinity for Lysozyme (Lyz) and Cytokeratin 20 (KRT20) respectively. We also found cells labelled solely by WGA, which may be early Paneth or goblet cells, and were referred to as immature mucus producing cells (IMPCs). Enterocytes were stained by Aldolase β (AldoB) as well as KRT20[2,6]. A number of cells were negative for all the used markers and were referred as transit-amplifying (TA) cells.

**Mapping endpoint cell types to lineages**. During time-lapse imaging, the Leica software allowed recording of the imaged locations, which could thus be found back after imaging. To achieve this, the mounting stage of the microscope and the orientation of the cover glass should be consistent with the settings during time-lapse imaging. With our optimized protocols for sample preparation and fixation (see Section Organoid fixation, permeabilization and staining), we could keep more than 50 % of the imaged organoids with limited deformations in the plate after fixation. After staining and relocating the organoids that were imaged in time during growth, mapping all cells (including their type information) to the cells that were tracked could be still challenging, due to the constant movement of cells during growth and global rotation and deformation of organoids caused by the fixation and repeated staining and washing. To mitigate these issues, we fixed the organoids within 5 minutes after the time-lapse imaging and performed every washing step gently, in order to preserve the spatial context of single cells. The linking of cells before and after fixation could be achieved mostly based on the spatial context of each cell. Linking was done in two steps. In a first automated step, we used a min-cost flow solver algorithm[7], which integrally optimizes the linking for all the tracked cells, and was also employed for the similar task of tracking cells between frames during organoid growth. In a second step, we manually corrected the automated linking results by visual inspection of the movie and staining images. The fluorescence intensity of H2B-mCherry showed heterogeneity between cells during time-lapse imaging, which could be preserved during fixation. Therefore, the brightness of the nuclear marker could also assist cell linking before and after fixation.

**Endpoint sister type analysis**. We studied the correlation of endpoint cell fates between sisters. For cells present at the endpoint, identified as specific type and with a sister, we checked the possible cell types of the sister pairs and counted the occurrence of each combination. For each cell type, we counted the number of sister pairs where at least one of them was of that type. If both sisters were of that type, the pair would be counted twice. Cells of different types had different abundance and majority of the sister pairs contained stem cells and/or TA cells. We then normalized the 2D histogram, via dividing the occurrence of each combination by the sum of each column, as shown in the bar plot in **Fig. 2a**. Therefore, the frequency within each column in **Fig. 2a** would sum up to be 1. The analysis was based on nine different organoids.

**Cell age distribution analysis**. For cells present at the endpoint, we could check the history of them and find the time when they were born. The duration between the birth time and the endpoint of imaging was measured as the cell age. Some cells were present from the beginning till the end. Their ages were then measured by the total length of the imaging experiment duration (~ 60 hours). The age distribution of each cell type was studied and plotted as a box plot in **Fig. 2b**, followed by statistical significance tests. Box plot elements represent the following: center line: median; box, quartiles; whiskers, range; fliers, outliers. The analysis was based on seven different organoids from experiments lasting ~ 60 hours.

**Analysis within (sub-)trees containing two cell types**. All of the (sub-) lineage trees with two different cell types were taken into account, unless more than 50 % of the cells within the lineage could not be tracked or died. The absolute count of each possible combination of the two cell types was shown in the 2D histogram (**Extended Data Fig. 6**).

**Cell type backpropagation**. The assumption underlying the backpropagation of cell types is that changes in cell types are rare. This assumption could be supported by the found type symmetry between sisters since frequent type changes likely lead to different types in sisters. Starting at the lineage endpoints, we propagate the measured endpoint types back in time following this process:

*__Backpropagation along consecutive timepoints.__ From one timepoint to a previous timepoint, the type is initially assigned as unchanged if no tree branch points (divisions) is traversed (marked '1' in __Fig. 2c__).*
*__Backpropagation of symmetric fate.__ For cell types identified by endpoint staining, we observed that sisters almost always assumed the same fate (__Fig. 2a__), suggesting that this fate was already set in the mother cell. Generalizing this observation, we assumed that if both sisters have the same (inferred) cell type, the inferred cell type of the mother cell is the same (marked '2' in __Fig. 2c__). Regarding cells with a dead sister, the mother is inferred the same type as the living daughter.*
*__Backpropagation of asymmetric fate__. The above backpropagation rule does not apply if two daughters have different (inferred) cell types. Therefore, we introduced two addition backpropagation rules. First, if at least one daughter's (inferred) cell type was stem cell, then the inferred cell type of the mother was also stem cell (marked '3' in __Fig. 2c__). Second, if the (inferred) cell type of one daughter was TA and the other daughter was not stem cell, then the inferred cell type of the mother was TA (__not shown in Fig. 2c__). These two rules were based on the capability of stem cells to generate all cell types and the transient property of TA cells between stem cells and differentiated cells.*
*__Forward propagation of cell type changes.__ If a mother and a daughter cell had different (inferred) cell types, we interpreted this as a change in cell type that occurred during the lifetime of the daughter cell (marked with triangle in __Fig. 2c,d__).*

These simple rules were sufficient to propagate backwards the lineage trees that we have encountered, unless the tree appeared very 'broken' where majority of the cells could not be tracked or died. All the lineage trees after backpropagation were shown in **Extended Data Fig. 4.**

**Imaging of a Lgr5 reporter**. To test our backpropagation method, we performed time-lapse imaging, endpoint staining and live-cell tracking in an organoid line with both Lgr5-GFP, a well-known stem cell marker, and a H2B-mCherry reporters. To limit phototoxicity caused by GFP imaging, we sampled the GFP channel about every 8 hours. The time-lapse imaging lasted for more than 24 hours, followed by endpoint staining.

Quantification of the membrane bound Lgr5-GFP fluorescence signals was conducted by determining the average fluorescence intensity within a 2D sphere with a diameter of 12 μm, which was sufficiently large to include one cell (**Extended Data Fig. 5a**). The Olfm4 staining was on the membrane as well and measured with the same method. The Lgr5 signal at the endpoint was plotted against the measured Olfm4 intensity (**Extended Data Fig. 5b**).

We quantified the fluorescence intensity of Lgr5-GFP in time. There were only few frames where Lgr5-GFP was imaged, therefore, most cells were only present in one or two of such frames. For cells only present in one frame, the measured GFP fluorescence within that frame would represent their Lgr5 signals. For cells present in multiple frames, their Lgr5 signals were obtained by averaging the GFP fluorescence between different frames. Quantification of Lgr5-GFP signals was performed for different (inferred) cell types (**Extended Data Fig. 5c**). For lineages inferred to lose stemness and transition to the TA type, we plotted the Lgr5 signals in time during the lineage progression (**Extended Data Fig. 5d**).

**Measuring locations of cells along the crypt-villus axis**. At each timepoint, the crypt-villus axis was manually annotated in the $xy$ plane at the $z$ position corresponding to the center of the crypt, since tracked crypts grew perpendicularly to the objective. Three to six points were marked along the axis, through which a spline curve was interpolated as the axis. For each tracked cell $i$ we determined its position along the spline by finding the value of $r_i$ that minimized the distance $d$ between the cell position and the axis (**Extended Data Fig. 7a**). The bottom-most cell of the crypt, i.e. that with the lowest value of $r_i$, was defined as position zero. Based on the shape and curvature of the epithelium, the location of crypt neck (where there was a sharp transition from crypt to villus) was estimated and annotated manually, as an indication of the length of the crypt. Since different crypts were of various length, we did a normalization of the locations based on the crypt neck location. For each cell's measured distance in μm within a certain frame, we divided it by the distance from the crypt neck within the same frame to the crypt bottom. Therefore, the length from crypt neck to bottom would remain one for each timepoint and each crypt. With this measurement, both the locations of different (inferred) cell types and the type transitions were mapped along the crypt-villus axis (**Fig 3.a-e**).

**Measurement of migration speed along the axis**. To estimate how fast a cell migrated along the axis, we searched for the locations of cells along the crypt-villus axis when they firstly showed up during tracking and the locations of cells when they were last present. The migration speed could be estimated by dividing the distance that the cell had migrated by the duration during which the cells were present (**Fig. 3f**).

**Search for neighbors for each cell**. Defining neighboring cells in organoids based on nuclear signal is non-trivial. Cells could have varying numbers of neighbors because of the disorder in the epithelium. Distances between nuclei could vary between cell types and location (spread apart in the villus-like region and closely packed in the crypts). To obtain robust neighbor pairs, we functionally defined neighbors as pairs of nuclei without another nucleus in between (**Extended Data Fig. 7d**). This condition was tested by a 'neighbor score', the ratio of the sum of the distances of the two cells of interest (A&B) to a third cell (S) and the distance between the two cells (A&B), namely $\frac{d_{AS}+d_{BS}}{d_{AB}}$. If the third cell S positioned perfectly in between the pair of interest A & B, the neighbor score would appear as the minimal value of 1 and A and B would not be identified as neighbors. If A and B were not separated by S, the three nuclei would form a triangle with high neighbor score between A and B. For each cell, we calculated the neighbor score for the twenty closest neighbors (in Euclidean distance) at every timepoint. If the neighbor score were higher than $\sqrt{2}$, we would consider them neighbors. This cut-off corresponded to diagonal neighbors in the case of a perfect square lattice. Using this cut-off, we found most cells with five or six neighbors, exactly as expected for the basal side of a curved epithelium[8].

**Measurement of separation rate**. Separation rates were determined by following pairs of neighbors over time. For a new born cell, its neighbors were searched and selected with the method introduced above. The selection was conducted one hour after division, so that the nuclei would have returned to the basal side of the epithelium. If the selected neighbors divided, we would continue tracking one of the daughters (selected randomly) so that the following of the neighbor pairs would not be cut short by division. The separation rates were measured after following the neighbor pairs for 2 hours and 10 hours, by calculating the fraction the pairs staying as direct neighbors within the total of pairs that were followed. Regarding the rearrangement rates of sisters, we followed the sister pairs that shared the same (inferred) cell type. Separation rates of sister pairs were also measured after 2 hours and 10 hours, by calculating the fraction of sisters staying as direct neighbors within the total number of sister pairs being followed (**Fig. 3h,i**).

**Analysis of isolated pairs of the same cell type**. We defined 'isolated pairs' as two nearby cells that are of the same type but surrounded by cells of other types. The two cells were either neighbors or one cell apart, therefore, they shared at least one common neighbor. Isolated pairs were identified at the endpoint using the introduced neighbor selection method (see Section Search for neighbors for each cell). We counted the occurrence of these pairs as sisters or not related cells and obtained the fraction of them as sisters. In this analysis, cells without a sister were excluded.

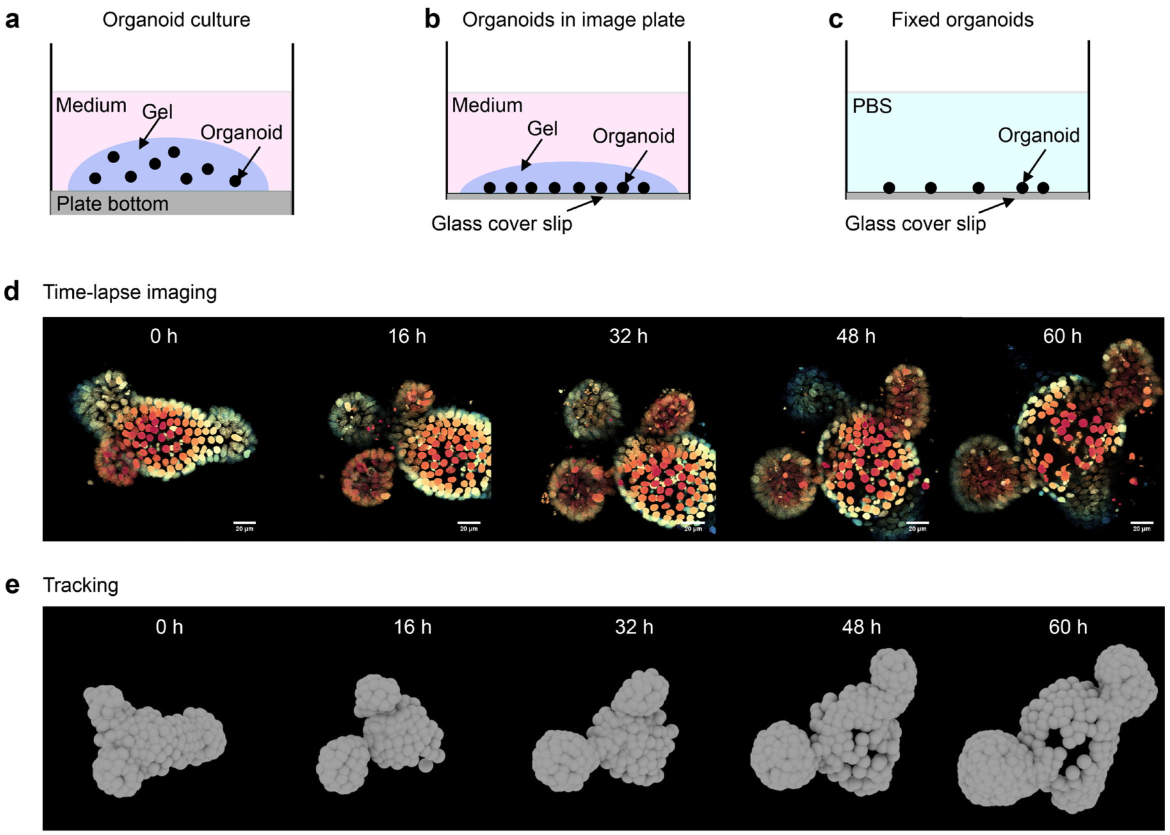


**f** The optimized staining order

| antibody binds to | round | excitation | emission | removal difficulty | staining signal |
|---|---|---|---|---|---|
| Olfm4 | 1 | 405nm | 421nm | Easy | Dim |
| CgA | 1 | 647nm | 675nm | Easy | Good |
| WGA* | 1 | 488nm | 518nm | Easy | Good |
| Aldoβ | 2 | 405nm | 421nm | Easy | Good |
| KRT20 | 2 | 647nm | 675nm | Difficult | Good |
| Lyz | 3 | 405nm | 421nm | Difficult | Good |

**Extended Data Fig. 1 | TypeTracker applied to mouse intestinal organoids. a,** For conventional organoid culture, organoids were scattered in ‘domes’ of BME gel and located at various heights. **b,** For time-lapse imaging, organoids were seeded in a thin layer of BME gel in chambered cover glass slides. Immediate after seeding, samples were put in fridges for 10 minutes so that organoids all sank towards the cover glass. **c,** With our optimized protocol, more than 50 % of the imaged organoids would remain at their imaged locations after fixation. **d,** Time-lapse imaging of an organoid carrying the H2B-mCherry reporter with 3D confocal for 60 hours. Scale bar, 20 μm. Color encodes different z planes. **e,** Live-cell tracking of the organoid. In these 3D reconstructions, each cell was represented by a sphere centered at the estimated nuclear center. Cells were not tracked if they were located far away from the objective and would move away from the region of interest. **e,** The order of antibodies and dyes to use in different rounds was optimized based on the staining quality and stripping difficulty of each antibody.

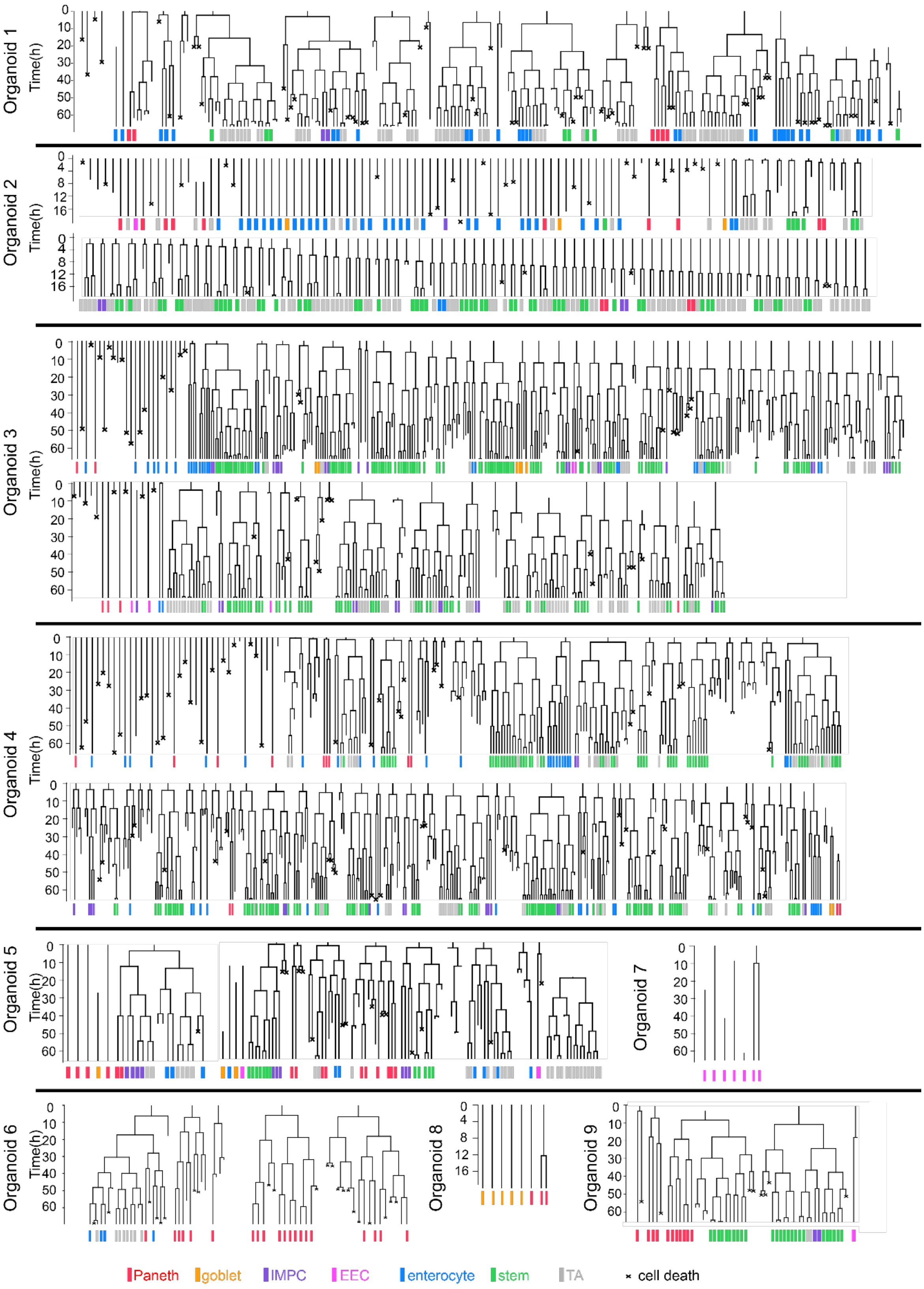


**Extended Data Fig. 2 | Gallery of lineage trees, generated from live-cell tracking, with cell types mapped at the endpoint.** These trees are from nine different organoids.

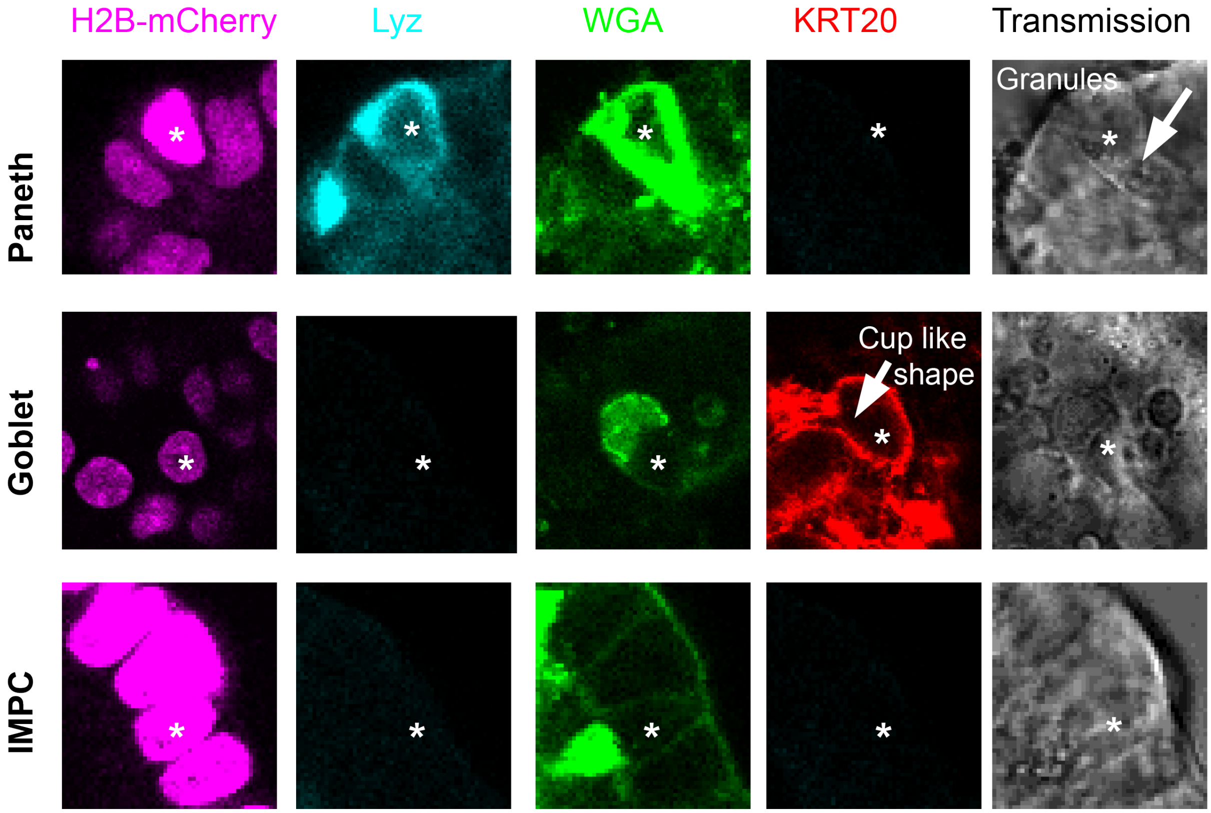


**Extended Data Fig. 3 | The identification of mature Paneth cells and goblet cells.** Paneth cells (indicated with * in row 1) often showed extremely bright H2B-mCherry fluorescence signals compared with the neighbor cells, bright Lysozyme (Lyz) fluorescence signals at the basal side of the cell, bright Wheat Germ Agglutinin (WGA) staining and granules in the transmission channel. Goblet cells (indicated with * in row 2) often stained positive of Cytokeratin 20 (KRT20) and WGA, with a cup like shape. A group of cells that stained positive of WGA but negative of KRT20 or Lyz were called Immature Mucus producing cells (IMPCs, indicated with * in row 3). These cells could be early Paneth cells or goblet cells considering the mucus secretion functions that they had.

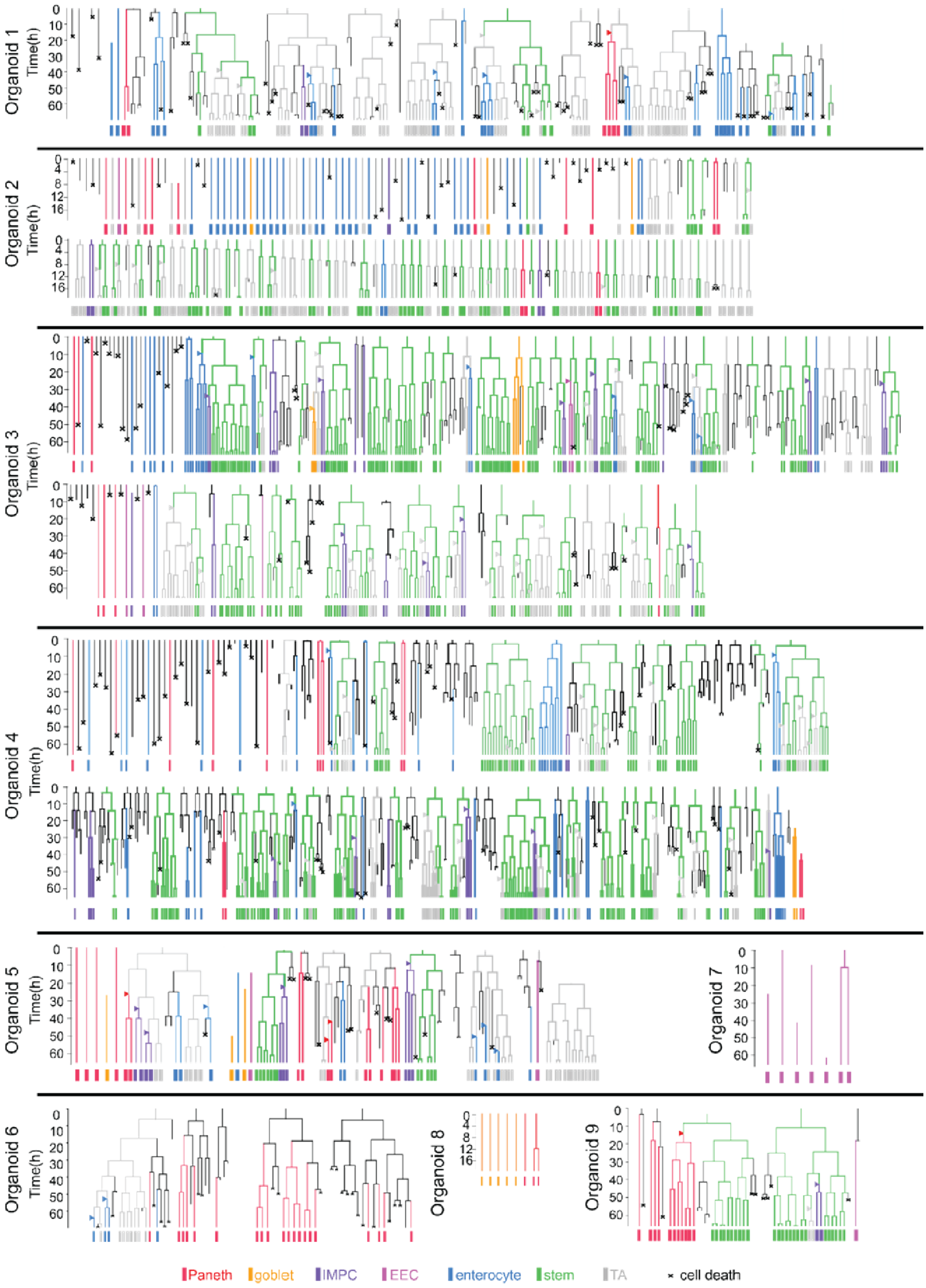

Organoid 1
Organoid 2
Organoid 3
Organoid 4
Organoid 5
Organoid 6
Organoid 7
Organoid 8
Organoid 9
Time(h)
Paneth
goblet
IMPC
EEC
enterocyte
stem
TA
cell death

**Extended Data Fig. 4 | Gallery of lineage trees after backpropagation with inferred cell type transitions.** Inferred cell types are shown with different colors. Type transitions are indicated by triangles, colored based on which type the cell was inferred to transition towards. These trees are from nine different organoids.

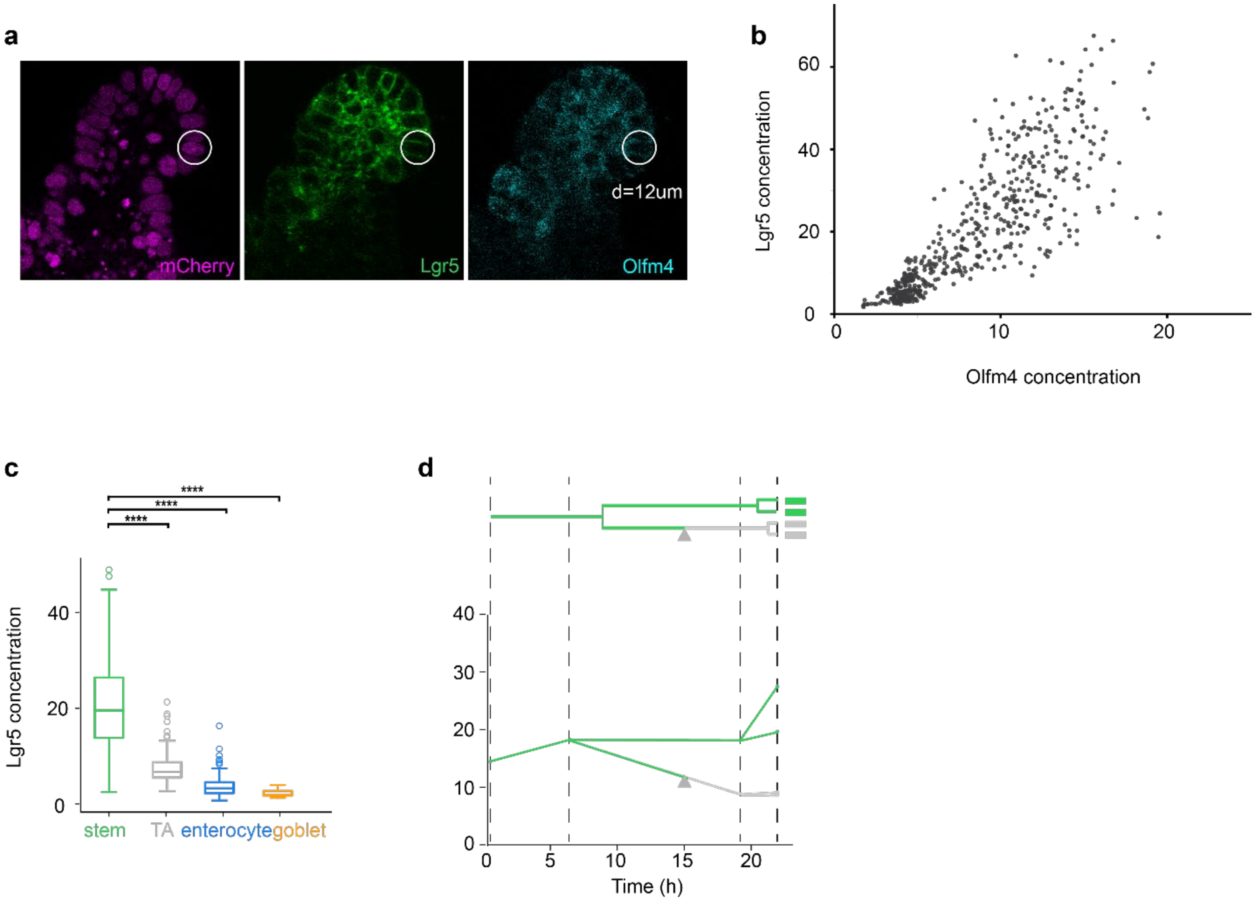


**Extended Data Fig. 5 | Imaging and tracking with a Lgr5 reporter confirmed the backpropagation method.** **a,** Images of the nuclei (H2B-mCherry), Lgr5-GFP (fluorescent reporter of stem cells) and Olfm4 (antibody used to identify stem cells). A 2D circle (d = 12 μm) was used to measure the fluorescence concentration of each cell. **b,** The Lgr5 and Olfm4 fluorescence concentration, measured by averaging the fluorescence intensity within 2D circles as shown in **a**, were proportional in single cells. **c,** The Lgr5 fluorescence concentration plotted against different (inferred) cell types, with stem cells showing high Lgr5 concentration, TA cells showing lower concentration and enterocytes and goblet cells having almost no Lgr5 fluorescence signals. **d**, Lgr5 fluorescence concentration measured in lineages going through transitions from stem cells to TA cells. The concentration measured ~ 4 hours after the inferred transition was much lower than the concentration measured ~ 8 hours before the transition.

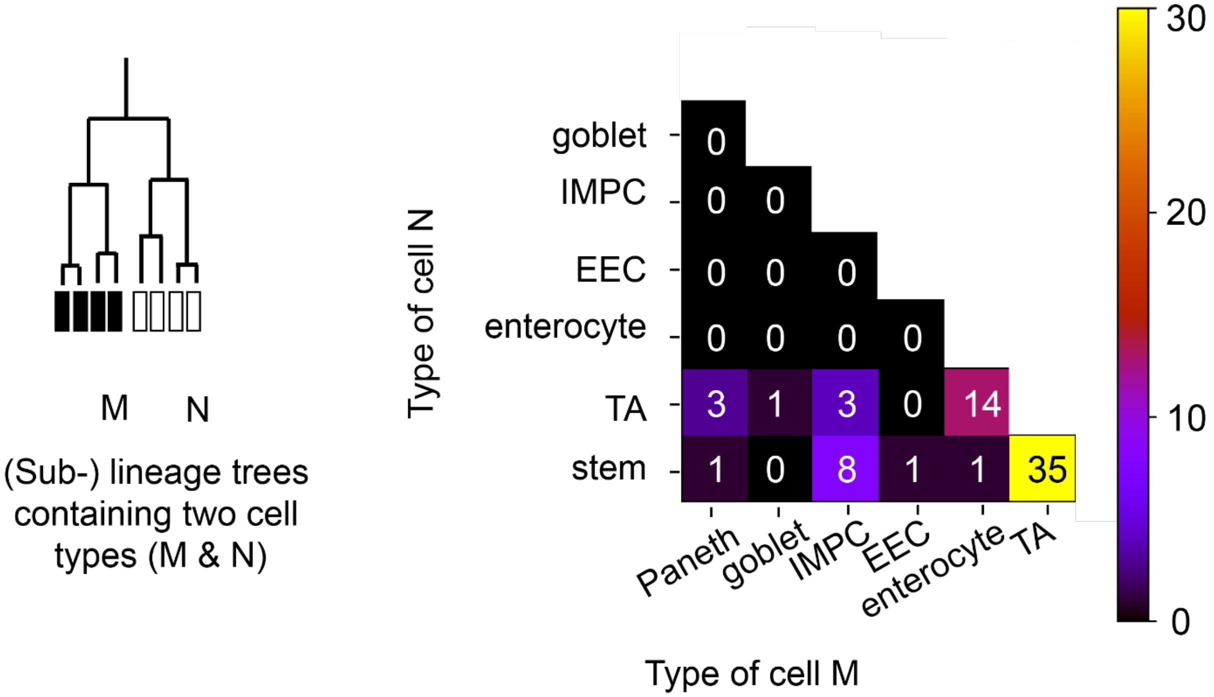


**Extended Data Fig. 6 | (Sub-) lineage trees containing two cell types often had at least one of the types as either stem cells or TA cells.** All of the (sub-) lineage trees with two different cell types were taken into account, unless more than 50 % of the cells within the lineage could not be tracked or died. The occurrence of each possible combination of the two cell types was counted and shown in the 2D histogram. Combination of two different differentiated cell types, such as enterocytes and goblet cells, was never found. Differentiated cells were often found together with either stem cells or TA cells in the same lineage.

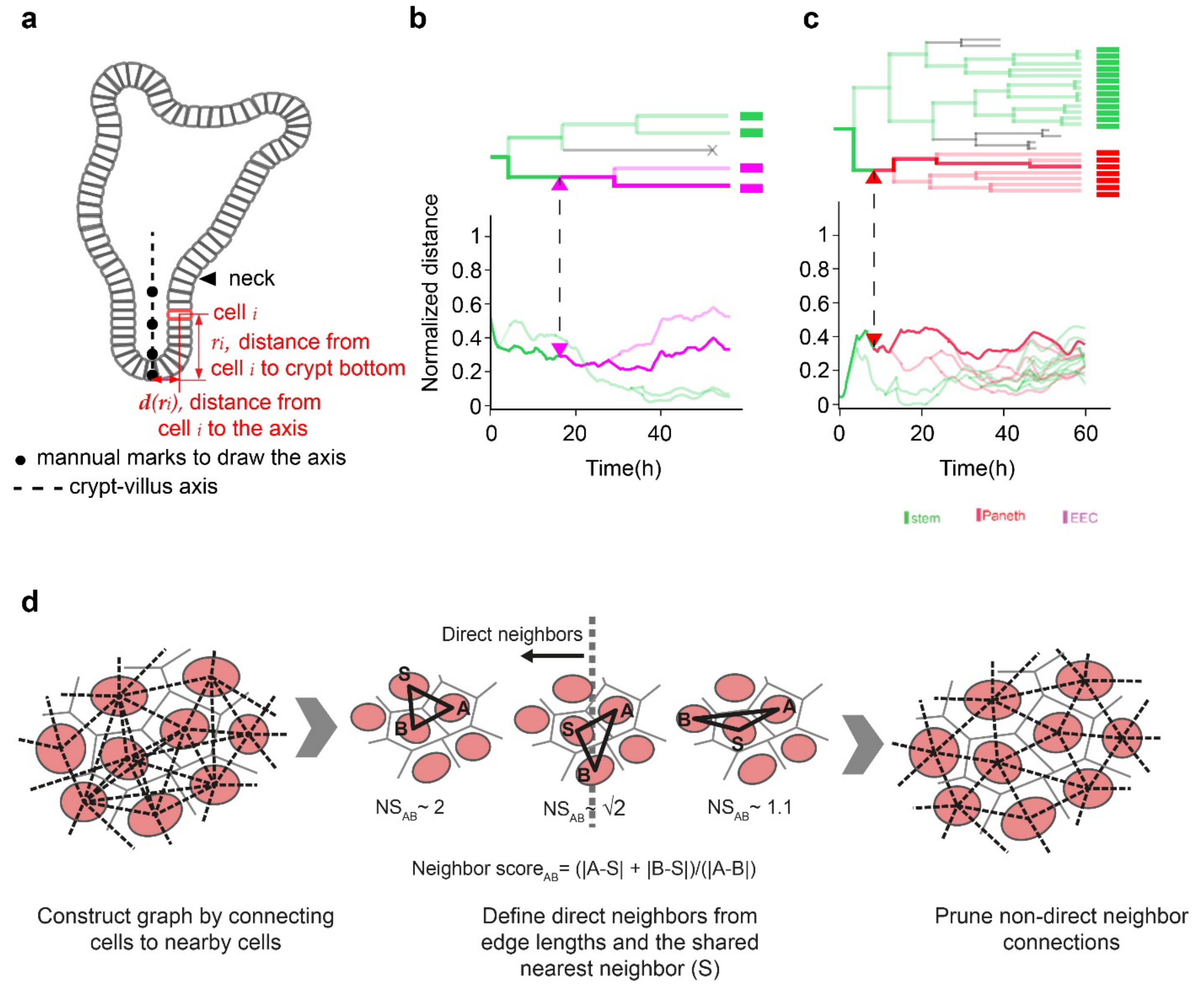


**Extended Data Fig. 7 | Following the spatial organization of cells during differentiation. a,** The crypt-villus axis could be generated by interpolating through the manually annotated points. For each tracked cell $i$, we determined its position along the axis by finding the value of $r_i$ that minimized the distance $d(r_i)$ between the cell position and the axis. **b & c,** The moving trajectories of cells within different lineages were colored by inferred cell types. Transitions to EECs and Paneth cells took place deep in the crypt, around 0.4, surrounded by stem cells which were often found from 0 to 0.6 along the axis. **d**, Neighbors were defined as pairs of nuclei without another nucleus in between. For each cell, the neighbor score for the twenty closest cells (in Euclidean distance) was calculated at every timepoint. If the neighbor score were higher than $\sqrt{2}$, cells would be identified as neighbors.
